# Nitro-Oleic Acid Induced Reactive Oxygen Species Formation and Plant Defense Signaling in Tomato Cell Suspensions

**DOI:** 10.1101/297994

**Authors:** Andrés Arruebarrena Di Palma, Luciano M. Di Fino, Sonia R. Salvatore, Juan Martín D’Ambrosio, Gustavo Esteban Gergoff Grozeff, Carlos García-Mata, Francisco J. Schopfer, Ana M. Laxalt

**Affiliations:** Instituto de Investigaciones Biológicas, CONICET-Universidad Nacional de Mar del Plata, Mar del Plata, Argentina; Department of Pharmacology & Chemical Biology, University of Pittsburgh, Pittsburgh, PA, USA; Instituto de Fisiología Vegetal CCT CONICET La Plata. Universidad Nacional de La Plata, Facultad de Ciencias Agrarias y Forestales. Diag. 113 N° 495, 1900 La Plata, Buenos Aires, Argentina

**Keywords:** nitro-oleic acid, tomato cell suspension, ROS, glutathione, signalling, plant defense

## Abstract

Nitrated fatty acids (NO_2_-FAs) are formed by the addition reaction of nitric oxide- and nitrite-derived nitrogen dioxide with unsaturated fatty acids. Nitrated fatty acids act as signaling molecules in mammals through the formation of covalent adducts with cellular thiols. The study of NO_2_-FAs in plant systems constitutes an interesting and emerging area. The presence of NO_2_-FA has been reported in olives, peas, rice and in Arabidopsis. To gain a better understanding of the role of NO_2_-FA on plant physiology, we analyzed the effects of exogenous application of nitro-oleic acid (NO_2_-OA) to tomato cell cultures. We found that NO_2_-OA induced reactive oxygen species (ROS) production in a dose-dependent manner via activation of NADPH oxidases, which requires calcium entry from the extracellular compartment and protein kinase activation, a mechanism that resembles the plant defense responses. NO_2_-OA-induced ROS production, expression of plant defense genes and led to cell death. The mechanism of action of NO_2_-OA involves a reduction in the glutathione cellular pool and covalently addition reactions with protein thiols and reduced glutathione. Altogether, these results indicate that NO_2_-OA triggers responses associated with plant defense, revealing its possible role as a signal molecule in biotic stress.

**Abbreviations:**

•NO_2_: nitrogen dioxide
•NO: nitric oxide
FA: fatty acid
GSH: reduced glutathione
H2O2: hydrogen peroxyde
NO_2_-FA: nitro fatty acids
NO_2_-Ln: nitro-linolenic acid
NO_2_-OA: nitro-oleic acid
OA: oleic acid
ROS: reactive oxygen species

## INTRODUCTION

Fatty acids (FA) not only provide structural integrity and energy for various metabolic processes to the plant cell but can also function as signal transduction mediators (Lim et al., 2017). As an example, oxylipins are oxygenated FAs, many of which are electrophilic species involved in plant defense against biotic and abiotic stresses (Lim et al., 2017; Farmer and Mueller, 2013).

Electrophilic nitro-fatty acids (NO_2_-FAs) are formed by the addition reaction of nitric oxide (•NO)- and nitrite (NO_2_^−^)-derived nitrogen dioxide (.NO_2_) to unsaturated fatty acids, in particular those containing conjugated double bonds (Schopfer et al., 2011; Baker et al., 2009). Electrophiles contain an electron-poor moiety, conferring attraction to electron-rich nucleophiles that donate electrons to form reversible covalent bonds via Michael additions (Chattaraj et al., 2006). In this regard, the electrophilic reactivity of nitroalkenes facilitates reversible addition reaction with cellular nucleophilic targets *(e.g*. protein Cys and His residues and reduced glutathione, GSH, Baker et al., 2007; Batthyany et al., 2006). This reactivity supports the post-translational modification of proteins, affecting their distribution and/or function. In addition, NO_2_-FA has been reported to act as •NO donors under certain conditions (Schopfer et al., 2005; Gorczynsk et al., 2007; Mata-Perez et al., 2016).

The study of NO_2_-FAs in plant systems constitutes an interesting and emerging area of investigation. The presence of nitroalkenes in plants was first reported in extra-virgin olive oil and linked to the beneficial effects of the Mediterranean diet on human health (Fazzari et al., 2014). In addition, NO_2_-FAs were later detected in Pea *(Pisum sativum)* and Rice *(Oryza sativa)* (Mata-Perez et al., 2017). Likewise, in cell suspensions of the model plant *Arabidopsis thaliana*, Mata-Perez et al., (2015) reported the presence of the nitroalkene nitro-linolenic acid (NO_2_-Ln). The levels of these NO_2_-FAs were modulated by both developmental stages and abiotic stresses (NaCl, low temperatures, cadmium or wounding). Moreover, treatments of Arabidopsis cell cultures with exogenous NO_2_-Ln induced differential gene expression related to oxidative stress responses as well as up-regulation of several heat shock response genes (Mata-Perez et al., 2015). In addition, in Arabidopsis roots and cell suspensions, NO_2_-Ln treatments induced •NO production (Mata-Perez et al., 2016).

Nitric oxide and reactive oxygen species (ROS) are signaling molecules involved in abiotic and biotic stress responses in plants. In this regard, tomato cell suspensions treated with pathogen-derived molecules, called elicitors like xylanase or chitosan displayed increased ROS and •NO production and induced plant-defense gene expression and cell death (Laxalt et al., 2007; Raho et al., 2011). During plant defense, NADPH oxidase activity of the Ca^2+^ and phosphorylation-dependent RBOHD (from respiratory burst oxidase homolog D) is upregulated, leading to increases in ROS production (Kadota et al., 2015). Thus, these physiological conditions where •NO and ROS are produced, provide a favorable chemical environment for the nitration of unsaturated fatty acids. Herein, we analyzed the signaling effects of exogenous treatment of tomato cell cultures with NO_2_-OA, with a particular focus on the induction of plant defense responses.

## RESULTS

### NO_2_-OA is Internalized and Metabolized in Tomato Cells

NO_2_-FAs are hydrophobic fatty acids with poor solubility in aqueous solutions. Thus, we first sought to analyze binding and internalization of NO_2_-OA by tomato cell suspensions. Figure 1A shows that NO_2_-OA effectively bound to tomato cells, reducing the remaining levels in media. Moreover, analysis of metabolic products of NO_2_-OA in treated cells revealed that NO_2_-OA is internalized and metabolized. In this regard, β-oxidation products and nitroalkene reduction products were detected (Figure 1B). These metabolites are a consequence of enzymatic reactions that take place in the cytoplasm and mitochondria of cells. These results indicate that NO_2_-OA is effectively internalized into the cell and therefore could be used as a model for the evaluation of NO_2_-FA physiological responses associated with its exogenous application.

**Figure 1.**
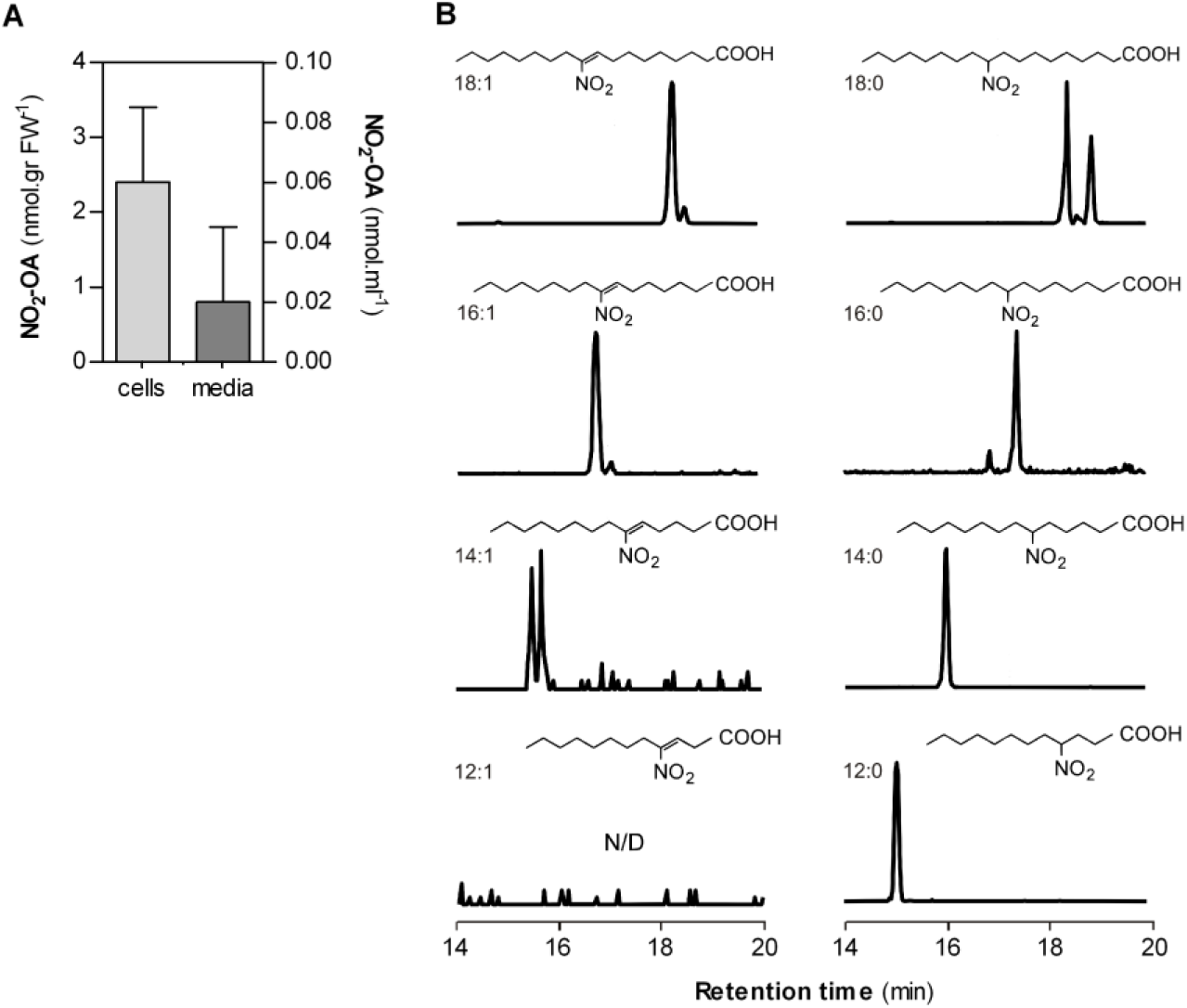
Detection and quantification of exogenous No_2_-OA added to tomato cell suspensions and metabolic products. Tomato cell suspensions were incubated with 10 µM No_2_-OA for 1 h and then No_2_-OA and its metabolise products were analyzed by HPLC-MSMS. A, Quantification of No_2_-OA in tomato cells or in the suspension media. Graph shows media with standard errors (n=3). B, Representative chromatographic profiles of No_2_-OA (left panel) and NO_2_-18:0 (right panel) and their β-oxidation products respectively found in tomato cells. N/D: not detected.

### NO_2_-OA Induces ROS but not •NO Production in Tomato Cells

Bioinformatics analisys of RNAseq data In Arabidopsis cell suspensions revealed that a large number of NO_2_-Ln-induced genes were related oxidative stress response, mainly depicted by hydrogen peroxide (H_2_O_2_) and reactive oxygen species (Mata-Perez et al., 2015). In this regard, we tested if NO_2_-OA could induce ROS production in tomato cell suspensions. As a control, we compared the response to oleic acid (OA), the non-nitrated backbone of NO_2_-OA. Figure 2A shows an increase in the fluorescence signal of NO_2_-OA-treated cells in a dose-dependent manner. Time course analysis showed that extending incubation times led to an increase in ROS production, with the exception of the 16 h treatment at 100 µM NO_2_-OA, where a decrease in ROS production was observed compared to 6 h. In the case of OA, none of the assayed conditions displayed any change in ROS production (Figure 2A). Fluorescence microscopy of tomato cells treated with 100 μM of NO_2_-OA for 6 h showed a significant increased in the fluorescent signal (Figure 2B).

**Figure 2.**
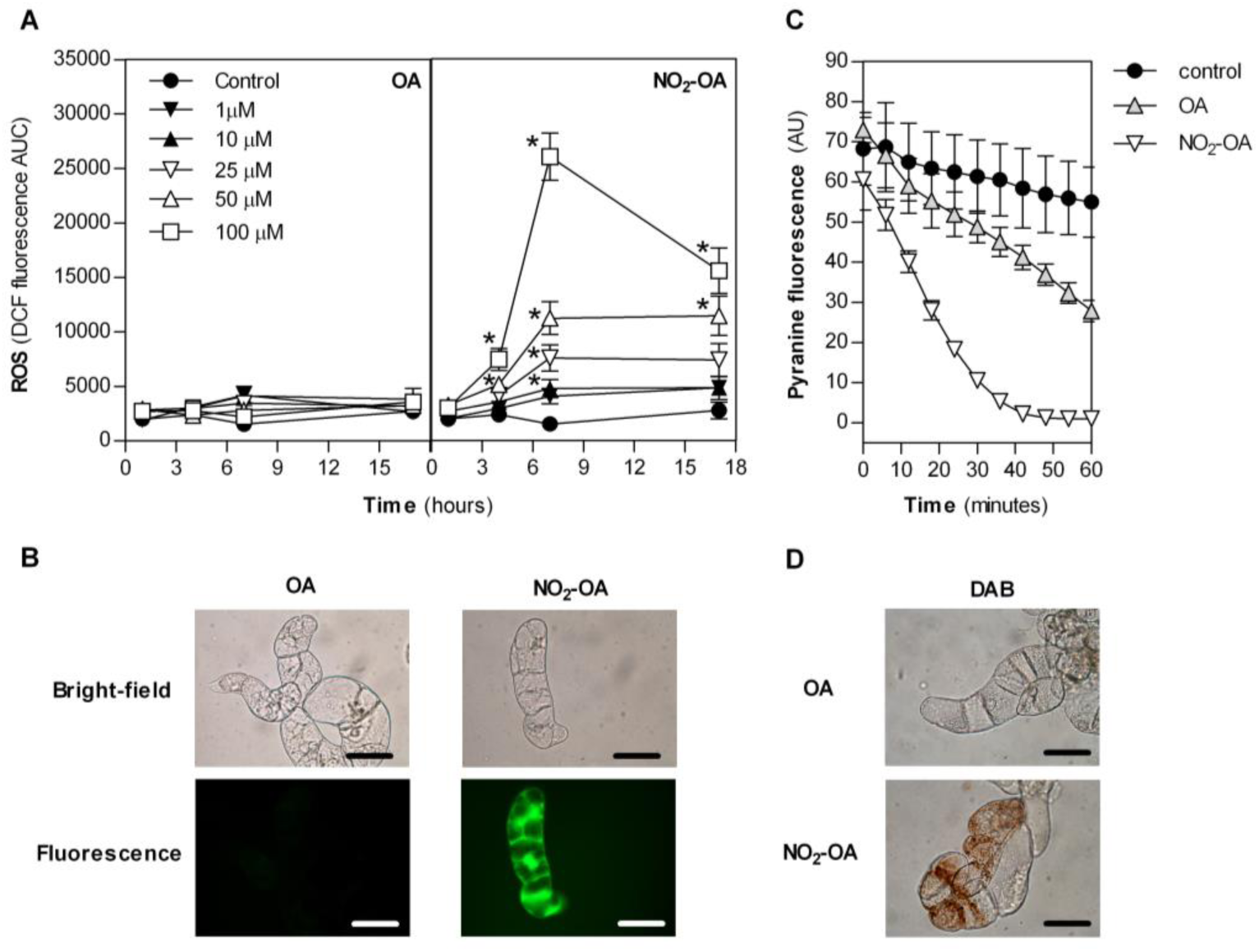
Reactive oxygen species (ROS) production in tomato cell suspensions treated with NO_2_-OA. A, Tomato cell suspensions were treated with OA or NO_2_-0A, or non-treated as a control. At 0, 3, 6 or 16 h of treatment 4 µM H_2_DCF-DA was added and the fluorescence was measured. The fluorescence was determined as the area under the curve (accumulated fluorescence within one hour). Data represents media and error standard of 4 independent experiments. * indicated significant difference (p<0.05) from control for each time (One way ANOVA post-hoc Holm-Sidak). B, ROS production on tomato cells suspensions treated for 6 h with 100 µM OA or NO_2_-OA and then incubated with 4 pM H_2_DCF-DA for 1 h. A representative light and epifluorescense microscope picture of experiments is shown. C, Oxidative burst. Cell suspensions were treated for 6 h with 100 µM OA or NO_2_-OA and then the quenching of pyranine fluorescence was recorded as a measure of the oxidative burst. Data represent media and error standard of 2 independent experiments. D, H_2_O_2_ detection by DAB stain on tomato cells treated with 100 µM OA or NO_2_-OA for 6h. Bars= 5 µm in panels B and D.

In order to further validate ROS production in NO_2_-OA-treated cells, we used two alternative methodologies. First, H_2_O_2_ production was analyzed using the pyranine quenching assay (Gonorazky et al., 2008). Figure 2C shows a rapid quenching of pyranine fluorescence in 100 μM NO_2_-OA-treated cells. To further confirm this increase in ROS, a second method based on 3,3′-diaminobenzidine (DAB) staining to detect H2O2 was used (Daudi and O’Brien, 2012). Again, NO_2_-OA treated cells showed positive staining with DAB when compared to OA-treated tomato cells (Figure 2D). Altogether these results show that NO_2_-OA but not OA triggers a dose- and time-dependent production of ROS in tomato cell suspensions.

Previous reports suggest that NO_2_-FA could act as a •NO donor in both, mammals and plants, a mechanism responsible for its physiological responses in cells (review in Baker et al., 2009, Mata-Perez et al., 2016). To test this hypothesis, tomato cells were treated for 1 and 6 h with NO_2_-OA and •NO production analyzed using the fluorescent probe DAF-FM-DA. NO_2_-OA was unable to induce •NO production in tomato cell suspensions at 1 h (data not shown) or 6 h of treatment (Supplemental Figure S2). These results indicate that under our experimental conditions NO_2_-OA does not act as a •NO donor and/or induce •NO production.

### NADPH Oxidase is Involved in NO_2_-OA-induced ROS Production

In plants, NADPH oxidase activation during plant defense is a key enzymatic source of ROS formation (Kadota et al., 2015). To specifically evaluate the role of NADPH oxidases as a source of ROS production triggered by NO_2_-OA, tomato cell suspensions were treated with the inhibitor diphenyleneiodonium (DPI). DPI treatments have been successfully used previously in cell suspensions and entire plant systems (Piedras et al., 1998; Govrin and Levine 2000; Orozco-Cårdenas et al., 2001; De Jong et al., 2004). In this regard, Figure 3 shows that addition of DPI to NO_2_-OA treated cells decreased ROS production in a dose-dependent manner.

**Figure 3.**
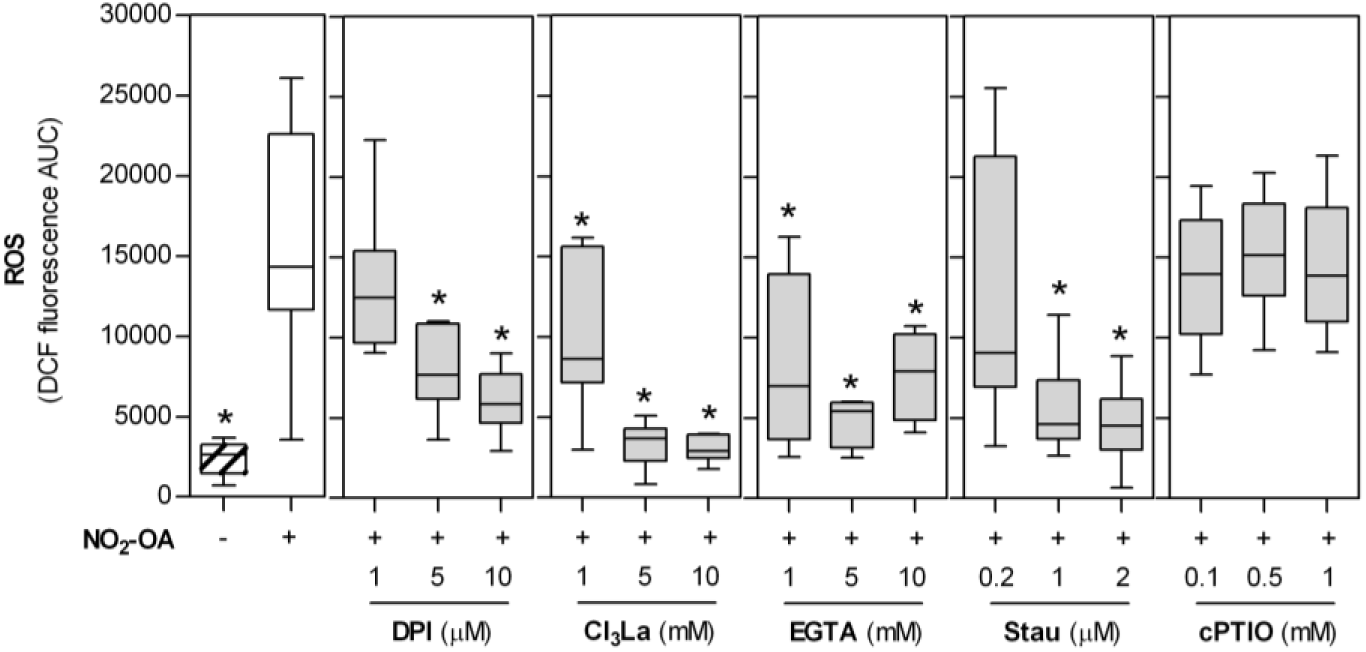
NO_2_-OA induced ROS production requires NADPH oxidase, Ca^2+^ and phosphorylation. Tomato cell suspensions were incubated with 100 µM NO_2_-OA for 6 hours (+) and as control, non-treated cells were incubated the same time (-). To 5 hours NO_2_^−^OA treated cells, different concentrations of NADPH oxidase inhibitor (DPI), calcium channel blocker (CI_3_La), extracellular calcium chelator (EGTA), protein kinase inhibitor staurosporine (Stau) or •NO scavenger (cPTIO) were added for another hour. Then, cells were incubated with 4 µM H_2_DCF-DAand the accumulated fluorescence was determined. Data is presented by box-plot were the box is bound by the 25th to 75th percentile, whiskers span to minimum and maximum values, and the line in the middle is de median of 6 experiments. * indicated significant difference from NO_2_-OA treated cells (One way ANOVA, post-hoc Holm-Sidak test, p <0.05).

NADPH oxidase-dependent ROS production is finely tuned by several signaling components, among them Ca^2+^, protein kinases and •NO-dependent posttranslational modifications (Kadota et al., 2015; Yun et al., 2011). Thus, we used a pharmacological experimental approach to assess the role of these signaling mechanisms on NO_2_-OA-induced ROS production. Both, the calcium channel blocker Cl_3_La and extracellular calcium chelator EGTA reduced ROS production triggered by NO_2_-OA (Figure 3). Thus, we conclude that ROS production in response to NO_2_-OA is triggered by Ca^2+^ entry from the extracellular compartment. Furthermore, the protein kinase inhibitor staurosporine decreased NO_2_-OA-induced ROS production (Figure 3) highlighting the requirement of phosphorylation events for the NO_2_-OA-dependent activation of NADPH oxidase. Finally, incubation of cells with the •NO scavenger cPTIO did not affect NO_2_-OA-induced ROS production (Figure 3). In aggregate, our results suggest that •NO is not involved in signaling responses leading to increased ROS formation elicited by NO_2_-OA in tomato cell suspensions.

### Induction of Plant Defense Gene Expression and Cell Death by NO_2_-OA

In tomato cells, we reported a rapid ROS production associated with the induction of gene expression and cell death upon treatments with the fungal elicitor xylanase (Laxalt et al., 2007; Gonorazky et al., 2014). Figure 4 shows the expression pattern of salicylic acid (SA)-dependent gene *SLPR1a*, a gene marker for hypersensitive response *SlHSR203J* and a jasmonic acid (JA)-dependent gene *SIPAL* at 3 h or 6 h upon treatment with NO_2_-OA or OA. No significant differences were found for any of the genes analyzed 3 h post treatment with NO_2_-OA. However, an increase in gene expression was observed for *SIPAL* and *SIPR1a* at 6 h.

**Figure 4.**
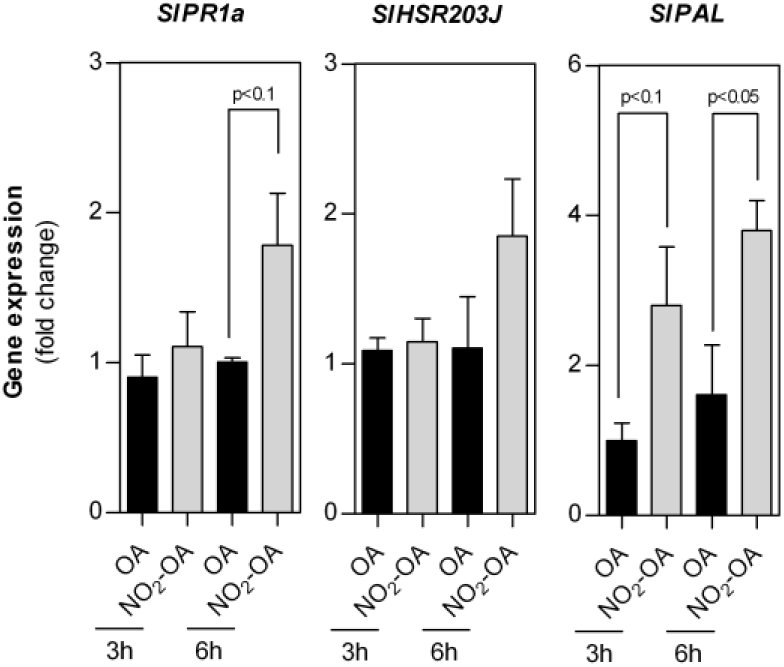
NO_2_-OA induces plant defence gene expression. Tomato cells suspensions were treated with 100 µM OA or NO_2_-OA. Cells were incubated for 3 or 6 h and total RNA was extracted. Transcripts levels of *SIPR1a, SIHSR203J* and *SIPAL* were analyzed by qPCR. *SIACT* (Actin) was used as a housekeeping gene. Data were analyzed by ΔΔCt method and fold change was calculated. Error bars represent standard deviations of media from 4 independent experiments. P values for each comparison are indicated in figure (One way ANOVA, post-hoc Holm-Sidak test).

The ROS burst and the increased expression of the above-analyzed genes suggest that NO_2_-OA could induce cell death. To evaluate the role of NO_2_-OA in this pathway, we determined cell death in tomato cells upon treatment with 50 or 100 µM NO_2_-OA or OA for 4, 7 and 17 h (Figure 5). Cells treated with NO_2_-OA at both tested concentrations lea to an increased rapid cell death rate compared to the corresponding OA treatment.

**Figure 5.**
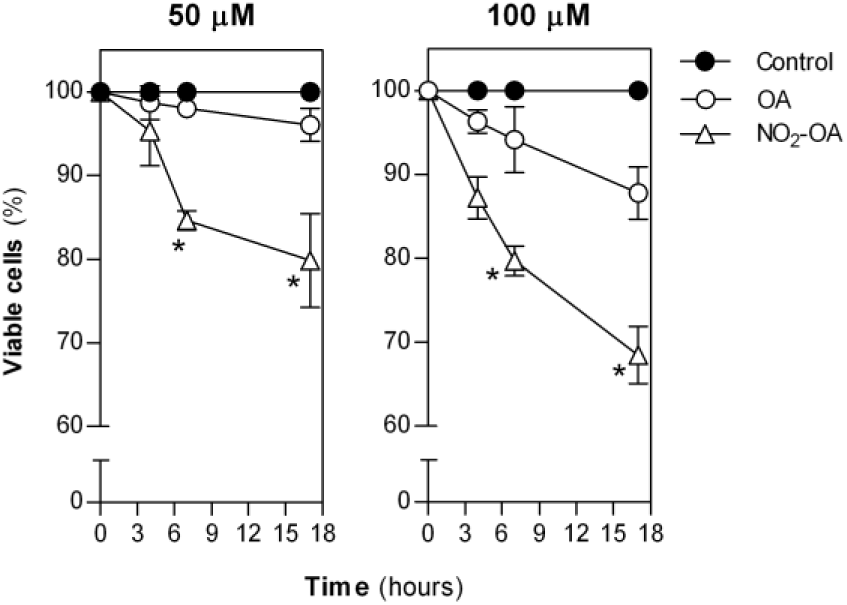
NO_2_-OA induces cell death. Tomato cells suspensions were incubated with OA or NO_2_-OA (at 50 µM or 100 µM) or without treatment as a control for 3. 6, and 17 h. Then, cells were stained with Evan Blue, incubated at room temperature for 5 minutes an observed under optical microscope. Live (none stained) and dead (blue stained) cells were manually counted on 10 random optical fields (40x). Data are expressed as percentage of viable cells respect to non-treated cells (black circles, control) at each time point. Data represents percentage average and standard error of 4 (0 and 3h) and 3 (6 and 17h) independent experiments. * indicated significant difference from control (z-test, p<0.05)

### NO_2_-OA Reduces Total GSH Content and Forms GS-NO_2_-OA and Protein-NO_2_-OA Adducts

NO_2_-FAs are electrophiles that can form adducts with several cellular nucleophiles, in particular with GSH and protein thiols (Freeman et al., 2008). Thus, we quantified the GSH pool (reduced and oxidized) in cells treated with 100 µM NO_2_-OA or OA for 3 h or 6 h to evaluate the extent of these reactions. Figure 6A shows that NO_2_-OA treatment led to a ~50 % decrease in total GSH. As this decrease was most likely associated with the formation of glutathione-NO_2_-OA adduct (GS-NO_2_-OA), we sought to detect their formation in tomato cells suspensions. In this regard, HPLC-MSMS analysis demonstrated the presence GS-NO_2_-OA adducts in NO_2_-OA treated cells (Figure 6B).

**Figure 6.**
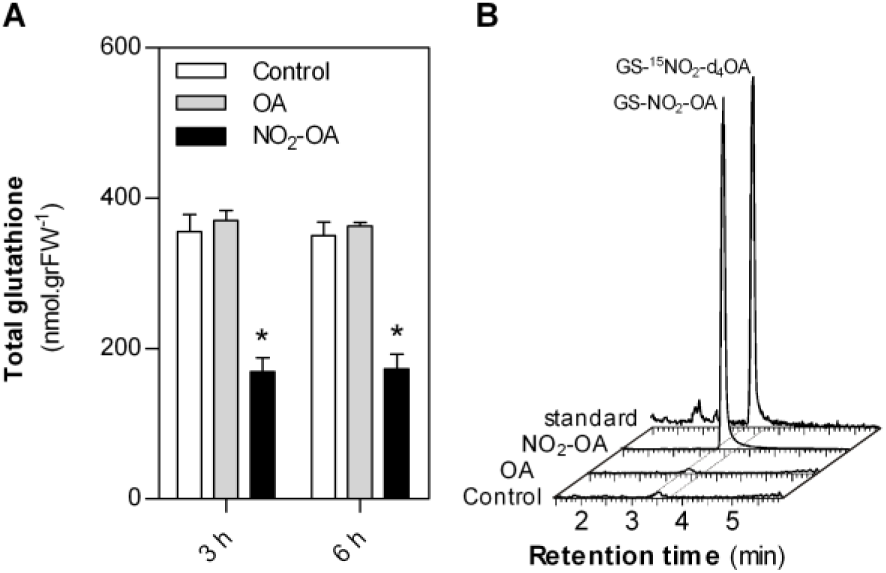
NO_2_-OA modifies glutathione cellular pool and forms GS-NO_2_-OA adducts. Tomato cell suspensions were treated with 100 µM NO_2_-OA or OA for 3 or 6 h. As control non treated cells were used. A, Total GSH pool was extracted and determinate by enzymatic GSH recycling method. Data represent media and standard error of 3 independent experiments. * indicated significant difference from control (One way ANOVA, post-hoc Holm-Sidak test, p <0.05). B, Detection of GS-NO_2_-OA adducts by HPLC-MSMS in tomato cell suspension treated with 100 pM OA or NO_2_-OA or without treatment for 3 h. Representative chromatograph form one of four independent experiments is show. As internal standard GS-^15^NO_2_-d_4_OA was used.

Given the detection of GS-NO_2_-OA adducts, we sought to evaluate the formation of protein-NO_2_-OA adducts in tomato cell suspensions. To this end, cells were incubated with NO_2_-OA conjugated to biotin for different times and the formation of protein-NO_2_-OA-biotin adducts was assessed at different times using western blot. Supplemental Figure S3 shows several tagged proteins in treated cells, indicating that cellular proteins are targets of NO_2_-OA. This further supports a role for protein covalent modification induced by NO_2_-OA in the signaling activities identified for this post-translational modification.

## DISCUSSION

Lipids function as signaling mediators in various plant processes with an important role in signal transduction. Signaling lipids in plants include a wide range of molecules such as glycerolipids, sphingolipids, fatty acids, oxylipins and sterols that participates in the response to different stresses like temperature, drought, wounding, nutrition starvation and pathogens among others (Wang, 2004). In this regard, NO_2_-FA represent a new class of lipid molecules involved in plant signaling. Sanchez-Calvo et al., (2013) proposed them to be novel mediators of NO dependent signaling pathways and metabolic processes in plant physiology. Later 9-NO_2_-cLA and 12-NO_2_-cLA isomers, were found for the first time in extra-virgin olive oil and NO_2_-OA was identified in whole olives adducted to cysteines (Cys-NO_2_-OA, Fazzari et al., 2014). In addition, NO_2_-Ln was detected in Pea, Rice and Arabidopsis. In the later, its levels changed during development and abiotic stress (Mata-Perez et al., 2015; Mata-Perez et al., 2017). Our attempts to detect free endogenous NO_2_-FAs in tomato cells suspension were unsuccessful. The source of plant fatty acids substrates to form nitroalkenes is an important aspect of these reactions that could involve membrane, mitochondrial and/or chloroplast phospholipids or triglycerides. In our experimental system, tomato cells are grown under dark conditions and have non-green plastids (Sello et al., 2017). Functional chloroplasts are very important for lipid signaling, particularly in defense responses to biotic stress (Serrano et al., 2016). In this sense, the fact that tomato cells have non-green plastids provides a plausible explanation for absence of NO_2_-FA in our measurements. We were unable to detected free NO_2_-FA in cells elicited with molecules derived from pathogens, such as xylanase, a condition that generates an oxidative and nitrosative stress (Laxalt et al., 2007) or during •NO donor treatments (data not show). However, when cells were pre-incubated with conjugated linoleic acid (cLA) and then treated with xylanase or •NO donors, cellular detection of NO_2_-cLA formation was observed (Supplemental Figure S4). This result indicates that tomato cells have the chemical environment required to endogenously nitrate fatty acids and generate electrophilic nitroalkenes. In humans cells, >99% of nitroalkenes are predicted to be covalently bound to thiols (Turell et al., 2017). The fact that we were unable to detect free NO_2_-FA could be due to the low levels of these nitro-lipids, their rapid metabolism, and/or the reversible chemical equilibrium established with thiols which favors adduct formation under cellular conditions. Given the uptake and metabolism of NO_2_-OA in tomato cell suspensions, we used it as a model system to study the effects of nitrolipids on plant defense responses.

NO_2_-OA induced ROS production in tomato cell suspension. This observation is in line with enhanced expression of several genes associated to H_2_O_2_ and ROS responses observed in Arabidopsis cell cultures (Mata-Perez et al. 2015). The inquiry of signaling downstream components of NO_2_-OA but upstream to ROS production, led us to find that calcium and phosphorylation events are required for ROS production. In plants, Ca^2^+ regulates ROS formation by NAPDH oxidase, through direct interaction with the Ct region of the protein, or by modulation of its activity through the action of CDPks (Kadota et al., 2015; Sagi and Fluhr 2006). Our results show that ROS production is independent of •NO, and occurs via activation of the NADPH oxidase, which requires Ca^2^+ and phosphorylation events. The presence of both signaling components in plant resembles the signaling pathway described in mammalian cells for NO_2_-FAs (Rudolph et al., 2010; Zhang et al., 2010).

ROS burst can lead to the up-regulation of several defense genes and cell death in tomato cell suspensions (Gonorazky et al., 2014). Particularly, we have previously demonstrated that upon xylanase treatment, there is an induction of plant-defense gene expression and cell death (Laxalt, et al., 2001; Laxalt et al., 2007). As mentioned above, in the presence of cLA, xylanase treatments provided the chemical environment required to generate electrophilic nitroalkenes. Exogenous addition of NO_2_-OA triggered the expression of defense response genes and cell death. Thus, under this condition, NO_2_-OA could be considered as a signaling component in plant immune response.

One mechanism of action of NO_2_-FAs involves their reactivity as electrophiles through Michael addition reactions with cellular thiols. We show evidence that NO_2_-OA modify the GSH cellular pool forming adducts with this NO_2_-FA. A similar response to sulforaphane, an electrophilic molecule was reported by Andersson et al., (2015) in Arabidopsis. Sulforaphane is a naturally occurring isothiocyanate derived from cruciferous vegetables that is present in widely consumed vegetables and has a particularly high concentration in broccoli. Sulforaphane reduced the GSH pool in Arabidopsis and increased cell leakage and cell death probably associated with ROS burst (Andersson et al., 2015). We determined that in tomato cells, sulforaphane induced ROS production in a similar way as NO_2_-OA does (Figure S5). Interestingly, as we demostrated for NO_2_-OA, sulforaphane can form adducts with cellular thiols thus generating post-translational modifications due to their electrophile nature (Groeger and Freeman, 2010). In summary, the post-translational modification of proteins and the GSH pool by Michael addition reactions of nitroalkene reveals a novel mechanism of action by which NO_2_-OA exert their activity in tomato cells. Future work will focus on the identification of protein targets adducted to NO_2_-FA. Altogether, we unravel the role of NO_2_-FA as a signal molecule in plant immune response.

## MATERIALS AND METHODS

### Tomato Cell Suspensions Culture Conditions

Tomato cell suspensions *(Solanum lycopersicum*, line Msk8) were grown at 25°C in dark in MS medium (Duchefa Biochemie, Haarlem, The Netherlands) as previously described (Laxalt et al., 2007). Cells of four-day-old cultures were used for all experiments.

### Chemicals and Reagents

OA was purchased from Nu-Chek Prep (Elysian, MN). NO_2_-OA and biotinylated NO_2_-OA were synthesized and purified as previously described (Woodcock el at., 2013; Bonacci et al., 2011; respectively). GS-^15^NO_2_-d4-OA standard was generated by the reaction of 200 mM reduced gluthatione with 100 µM ^15^NO_2_-d_4_-OA in 50 mM phosphate buffer (pH 8 at 37°C) for 3 h. The lipid conjugates were loaded on a C18 SPE column pre-equilibrated with 10% methanol and then eluted with methanol. Solvents used for extractions and mass spectrometric analyses were of HPLC grade or higher from Burdick and Jackson (Muskegon, MI).

### Lipid Extraction

Lipid extraction from 100 mg of tomato cells were carried out using hexane:isopropanol:1M formic acid (2:1:0.1, v/v/v). As internal standard samples were spike with ^15^NO_2_-d_4_-OA (100 nM). The organic phase was dried under N_2_ and reconstituted in methanol before MS analysis.

### Chromatography

Nitro-FA and GS-NO**2**-OA were analyzed by HPLC-ESI-MS/MS using gradient solvent systems consisting of water containing 0.1% acetic acid (solvent A) and acetonitrile containing 0.1% acetic acid (solvent B), and were resolved using a reverse phase HPLC column (100 × 2 mm × 5 µm C18 Luna column; Phenomenex) at a 0.65 ml/min flow rate. NO**_2_**-FA samples were applied to the column at 30% B (0.3min) and eluted with a linear increase in solvent B (100% B in 14.7min) and GSH adducts were applied to the column at 20% B (1.1 min) and eluted with a linear increase in solvent B (20-100% solvent B in 5.9 min).

### Mass Spectrometry

The NO**_2_**-FA detection was performed using multiple reactions monitoring (MRM) on an AB5000 triple quadrupole mass spectrometer (Applied Biosystems, San Jose, CA) equipped with an electrospray ionization source. MS analyses for NO**2**-FA used electrospray ionization in the negative ion mode with the collision gas set at 4 units, curtain gas 40, ion source gas #1 55 and #260, ion spray voltage -4500 V, and temperature 600 °C. The declustering potential was -100, entrance potential -5, collision energy -35, and the collision exit potential -18.4. MRM was used for sample analysis of nitrated fatty acids following the charged loss of a nitro group (m/z 46) upon collision-induced dissociation. An AB6500+ Q-trap triple quadrupole mass spectrometer (Applied Biosystems, San Jose, CA) was used for GSH adducts detection in positive ion mode using the following parameters: electrospray voltage 5.5 kV, declustering potential 60 eV, collision energy 30, gas1 45 and gas2 50 and de source temperature was set at 550°C. The following transitions 635.2/506.2 and 640.2/511.2 were used for detecting GS-NO_2_-OA and GS-^15^NO_2_-d4-OA respectively.

### Determination of ROS and •\NO Production

Tomato cells (90 µL per well in 96-well microtitre plate, DeltaLab) were treated with 1, 10, 25, 50 or 100 µM of OA or NO_2_-OA for 1, 4, 7 or 17 h. Plates were incubated at 25°C in darkness. ROS production was detected by incubating cells with 4 μM H_2_DCF-DA probe (Ubezio and Civoli, 1994; Molecular Probe, Eugene, OR, USA) during the last hour of each treatment. As an example, for 7 h treatment, at 6 h 4 µM of H_2_DCF-DA was added and ROS production was measured as follow. Cells were immediately introduced in Fluoroskan Acsent microwell fluorometer (Thermo Electron Company, Vantaa, Finland) and fluorescence (ex 485nm, em 525nm) was recorded every 2 minutes for 60 minutes. The area under the curve (AUC, accumulated fluorescence) was calculated according to equation showed in supplemental data and taken as an accumulated florescence value (see supplemental Figure 1S). For •NO determination 10 μM DAF-FM-DA was used as a probe (Kojima et al., 1999, Molecular Probe, Eugene, OR, USA) and production was calculated as indicated above for H_2_DCF-DA.

For observation of ROS production, cells were treated with 100 μM of OA or NO_2_-OA for 6 h and then incubated with H_2_DCF-DA for 1 h and visualized under the epifluorescence microscopy with an excitation filter of 495 nm and a barrier filter of 515 nm according to Gonorazky et al., (2008).

Hydrogen peroxide determination was carried out by Pyranine quenching assay according to Gonorazky et al., (2008, Pyranine Sigma-Aldrich, St. Louis, MO, USA). Fluorescence quenching was recorded every 2 minutes for 60 minutes using Fluoroskan Acsent microwell fluorometer.

*In situ* hydrogen peroxide production was assayed by DAB staining. Briefly, 100 µl of treated cells were incubated with 50 µl of 0.2% DAB solution (Sigma-Aldrich) prepared according to Daudi and O’Brien, (2012). Cells were incubated over night and observed under microscope.

### Inhibition Assays of ROS Production

Tomato cell culture were treated in 96-well microtitre plate (90 µL per well) for 5 h with 100 µM of NO_2_-OA and then incubated with different concentrations of NADPH oxidase inhibitor (DPI: 1, 5 or 10 µM, Sigma), calcium channel blocker (Cl_3_La: 1, 5 or 10 mM, Sigma-Aldrich), extracellular calcium chelator (EGTA: 1, 5 or 10 mM, Sigma-Aldrich), protein kinase inhibitor (staurosporine: 0.2, 1 or 2 µM, Sigma-Aldrich) or •NO scavenger (cPTIO: 0.1, 0.5 or 1 mM, Invitrogene, Carlsbad, CA, USA) for an additional hour in presence of 4 µM H_2_DCF-DA. Control cells (no treatment, negative control) and NO_2_-OA-only treated cells (positive control) were incubated under the same conditions. Determination of ROS production was performed as indicated above.

### qPCR Analysis of Gene Expression

Three ml of tomato cells cultures were treated with 100 µM OA, 100 µM NO_2_-OA or DMSO (Merk, Darmstadt, Germany) as a control for 3 or 6 h. Cells were washed with phosphate buffer (pH 7.5, 50 mM), frozen in liquid nitrogen and total RNA was extracted using the Trizol method. cDNA was synthesized according to manufactured instruction using M-MLV enzyme (Invitrogene). Transcripts levels of *SlPR1a, SIHSR203J, SlPAL*, and *SIACT* (Actin) genes were analyzed by qPCR (StepOne, Thermo). Expression data are expressed as ΔΔC_t_ and *SIACT* was used a housekeeping gene. Primers used are listed in supplemental Table S1.

### Cell Death Quantification

Tomato cells were treated with 50 µM or 100 µM of OA or NO_2_-OA for 4, 7 or 17 h on 96-well microtitre plate (90 µL per well). At each time, 50 µl of 1% ^w^/v Evans Blue solution (Fluka, Buchs, Switzerland) were added to cells in wells, incubated at room temperature for 5 minutes and observed under light microscope. Live (none stained) and dead (blue stained) cells were manually counted on at least 10 random optical fields (40x) for each treatment.

### GSH and GS-NO_2_-OA Adduct Detection

Three ml of tomato cell culture were treated with 100 µM OA, 100 µM NO_2_-OA or DMSO as control for 3 or 6 h. Cells were collected, washed and immediately frozen in liquid nitrogen. Total GSH was evaluated using the enzymatic GSH recycling method (Griffith, 1980).

GS-NO_2_-OA adducts were assessed by HPLC-MSMS. A mass of 0.4 mg of cell was spike with 30 fmol of GS-^15^NO_2_-d_4_-OA as internal standard before extraction. GSH adducts was extracted using C18 SPE columns. Columns were conditioned with 100% methanol, followed by 2 column volumes of 10% methanol. Samples were loaded into the SPE column and washed with 2 column volumes of 10% methanol and the column was dried under vacuum for 30 min. GSH adducts were eluted with 3 ml methanol, solvent was evaporated, and samples were dissolved in methanol for analysis by HPLC-electrospray ionization mass spectrometry (ESI-MS/MS).

### Western Blot of Protein-NO_2_-OA Adducts

Tomato cell cultures (500 µl) were treated with NO_2_-OA-biotin at a final concentration of 25 μM for 4, 7, and 17 h. As a control, 500 μl of cell cultures were treated with DMSO. The cells were collected, subjected to three cycles of freeze/thawed and ground under liquid nitrogen for mechanical disruption. Proteins were extracted using phosphate buffer (50 mM pH 7.5) containing 20 mM NEM (Fluka). Total protein concentration was determined by the bicinchoninic acid method (Smith et al., 1985, bicinchoninic Sigma) and 100 µg of proteins for each sample were reduced by incubation with 10 mM BME (BioBasic, Ontario, Canada) for 5 minutes at 70°C (Schopfer et al., 2009). As a positive control, 100 µg of tomato proteins cell extract were treated with an excess of NO_2_-OA-biotin (125 µM final concentration) to induced nitroalkylation (room temperature for 30 minutes in phosphate buffer). Samples were treated with BME and heat as indicated above. All samples were mixed with protein loading buffer without BME, separated in polyacrylamide gels, transferred to nitrocellulose membrane and incubated with mouse anti-biotin primary antibody overnight (Sigma-Aldrich). The membrane was incubated with a secondary antibody coupled to phosphatase alkaline enzyme (Sigma-Aldrich) for 3 h and developed over 5 minutes or 2 h (see supplemental Figure S3).

